# Lack of evidence for the presence of an interferon in invertebrate

**DOI:** 10.1101/000513

**Authors:** Pei-Hui Wang

**Author notes:** Address correspondence and reprint request to Pei-Hui Wang, E-mail address (P.-H. Wang).

## Abstract

In vertebrates, the interferon (IFN) response is the primary form of innate antiviral defense. Previously (2005), a partial cDNA which could encode an interferon-like protein (IntlP) is reported in shrimp, later Rosa et al. (2008) argue that this partial cDNA should encode a portion of insect mitochondrial ATP synthase (MAS) B-chain. Recently (2009), it is demonstrated IntlP also possess antibacterial activity beside antiviral activity reported before. Lacking of a consensus opinion to the question of whether this gene encodes IntlP or MAS, we try to provide more evidences to identify this gene exactly. Here we obtain the full length cDNAs of IntlP/ MAS in *Litopenaeus vannamei*, and perform the tissue distribution and induced expression analysis. Our results confirm that IntlP is coded by a mistaken ORF and the actual protein indeed is a *L. vannamei* mitochondrial ATP synthase (LvMAS) whose function is unknown in antiviral responses.

## 1. Introduction

Interferons (IFNs) constitute a large group of cytokines that are best known for their ability to induce vertebrate cells into an antiviral state. It is also reported IFN system can defense against bacterial and protozoal infection. Binding of IFNs to their receptors initiates signaling that leads to a global shutdown in protein translation, cellular RNA degradation and deamination and often the death of virus-infected cells. However, until recently, there was no IFN cDNA of invertebrates found. In 2005 He et al. reported a partial cDNA encodes an interferon-like protein (IntlP) homologue to mammalian IFN-αwhich was expressed only in the WSSV-resistant shrimp *Penaeus japonicus* (but not in naïve shrimps) and showed non-specific antiviral activity to SGIV (grouper iridovirus). But later Rosa et al. (2008) argue that this partial cDNA actual encodes a portion of the mitochondrial ATP synthase (MAS) which shows high identity (60–73%) with insect MAS b-chain and was expressed not only in naïve and WSSV-infected *L. vannamei* but also in two wild Brazilian shrimp species. As well as He, Rosa didn’t obtain the full length cDNA of shrimp ATP synthase. Thus, like IntlP, it is unclear whether this portion of the ATP synthase is in the right coding region of the full length cDNA. It is also unknown whether this gene has an induced expression by pathogen infection and where it localizes (secreted or not) for function. Recently, Mai et al. (2009) report IntlP also possesses significant antibacterial activity to the shrimp pathogens *V. alginolyticus* and *V. parahemolyticus*. About this partial cDNA encoding IntlP or MFS has two different points of view. Maybe more empirical evidences are needed to confirm a substantive protein encoded by this gene. We obtain the full length cDNAs of IntlP/MAS in *L. vannamei* by RACE-PCR approach, and then several cDNAs show very high nucleotide identities (>60%) with *L. vannamei* MAS (LvMAS) in other lobsters and crabs were retrieved in NCBI. All these cDNA sequences encode proteins show high identities with insect MAS but very low with mammal IFNs. RT-PCR reveals that LvMAS mRNA cannot be induced by immune challenge. Further homology searching and sequence analysis indicate IFNs most probably origin from cartilaginous fish and to date no correct invertebrate IFN cDNAs have been cloned. This study also makes it clear that IFN system is limited in high vertebrates, while RNA interference is used by nematode and insects as a main antiviral strategy. And this also points an interesting question that which mechanism is used in the ancestor of high vertebrate like sea urchins, amphioxus, hagfish, and lamprey and so on which lack IFN system and no RNA interference pathway found.

## 2. Materials and methods

Based on the partial cDNA (accession no. **<u>EU246975</u>**) which could encode an interferon-like protein (IntlP) in *L. vannamei*, specific primers (Table. 1) were designed to obtain the 3’ and 5’ end cDNA sequences of LvMAS by rapid amplification of cDNA ends (RACE) as described before (Wang et al., 2009). The genes of American Lobster, Blue Crab, *Petrolisthes cinctipes* and Grass Shrimp show high nucleotide identities (﹥ 70%) with LvMAS were obtained through the NCBI programs blastn and blastx, and the ORFs were predicted through the NCBI program ORF Finder. RT-PCR was performed with LvMAS-F and LvMAS-R. cDNA templates for RT-PCR were prepared previously (Wang et al., 2009), and the conditions were the same as described before except that the cycles were modified as indicated in Fig. 2B. Reported vertebrate IFN-α and IFN-γ were used as seed sequences to search IFN homologous of *C. elegan*, insects, sea urchin *S. purpuratus*, amphioxus, lamprey in UCSC Genome Browser and NCBI using “BLAST”. When search IFN homologous of pacific oyster *Crassostrea gigas*, shrimps and crabs, databases of Marine Genomics Project (http://www.marinegenomics.org/) are also referred using the provided search tool.

**Table 1.**
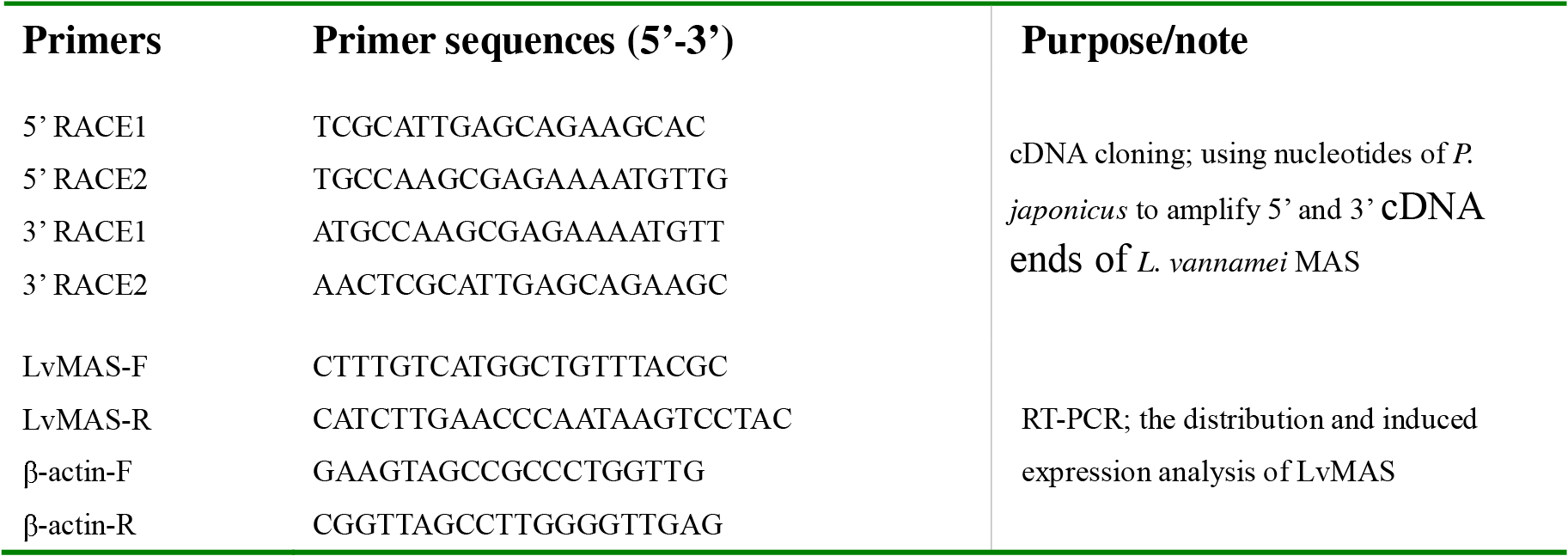
PCR primers used in this study.

## 3. Results and discussion

The predicted ORFs of American Lobster, Blue Crab, *L. vannamei*, Petrolisthes cinctipes and Grass Shrimp all encode a protein (AlMAS, BcMAS, LvMAS, PcMAS and GsMAS) which possesses a mitochondrial ATP synthase domain; no alternative ORF which could encode a protein possesses an interferon domain is available (Fig. 1D). The ORF encodes the IntlP used by He et al which does not contain an interferon domain and shows very low identity with vertebrate IFN-α (Fig. 1B). As later Rosa described, the analysis of IntIP gene through the NCBI programs blastx resulted in a translated nucleotide sequence that strongly matched with the MASs of other species (Fig. 1C). In addition, IFNs are secreted proteins which possess a signal peptide in the N-terminus. But we can not find any signal peptide using all possible translation patterns of these five genes (Fig. 1D). The obtaining of the full length cDNA of LvMAS and further sequence analysis make us believe that IntlP most probable is encoded by a mistaken ORF. To confirm this conclusion, RT-PCR was performed to investigate the distribution and induced expression of LvMAS (Fig. 2B). We observe that LvMAS is wildly distributed in healthy and immune challenged shrimp *L. vannamei*, a result correlated with later Rosa. When challenged by saline, LPS (from *E. coli*), Gram-negative *V. alginolyticus*, Gram-positive *S. aureus*, Yeast *S. cerevisiae*, white spot syndrom virus (WSSV) or polyinosinic polycytidylic acid (poly I: C) as described before, the expression of LvMAS has no obvious changing (Fig. 2B). The constitutive expression of LvMAS is more like MAS rather than IFNs which could be highly induced after immune challenge. As for the results that recombinant IntIP possess non-specific antiviral and antibacterial activities described by He et al and Mai et al, it remains unexplained. In the article (He et al., 2005), Fig. 3B displays the antiviral activity of recombinant IntlP by a cytotoxicity experiment by inhibiting SGIV on fish GP cell lines (grouper embryo cells). According to the authors, GP cells (Fig. 3A) were completely destroyed by SGIV (Fig. 3B) while parts of the cells remained alive when incubated in the SGIV and IntlP protein mixture for 48 h, (Fig. 3C). But the fingers are not clear enough and no parallel experiments were declared. In the paper (Mai et al., 2009), three different methods are mentioned in detecting IntlP antibacterial experiments, but they were only done in duplicates and no significant differences were calculated. And in these two papers, the antiviral and antibacterial assays both lack positive control groups. In the later paper (Mai et al., 2009) in Table 1, a negative group is also lacking. In addition, IntIP indeed shows some similarity with a portion of mammalian IFNs although it is very low, so it can not be rule out that recombined IntIP protein holds some functions like the IFNs.

**Fig.1.**
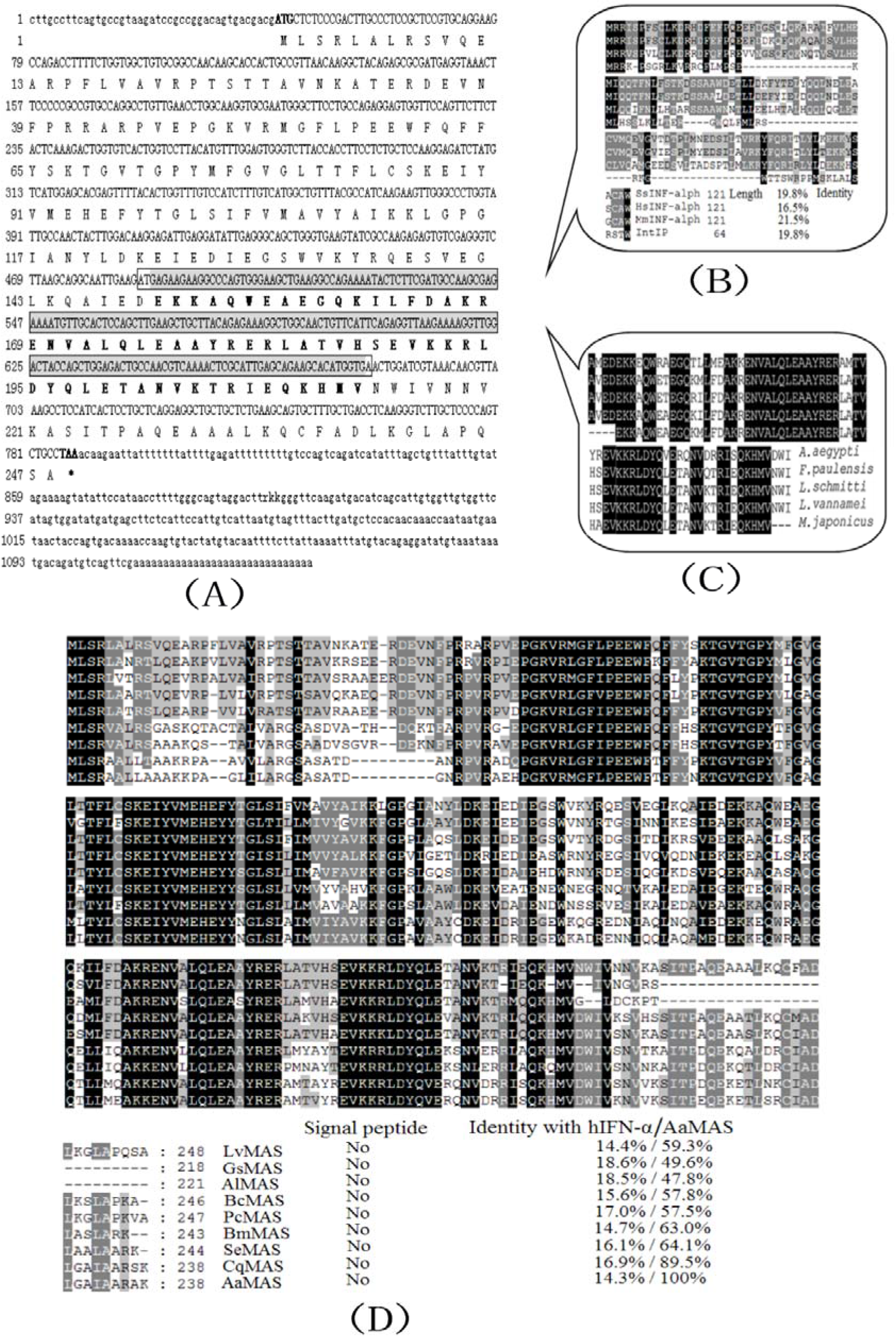
cDNA cloning (A) and sequence analysis (D) of *LvMAS*. The ORF encodes the IntlP used by He et al shows very low identity with vertebrate IFN-α(B). The analysis of IntIP gene through the NCBI programs blastx resulted in a translated nucleotide sequence that strongly matched with the MASs of other species (Fig. 1C).

**Fig.2.**
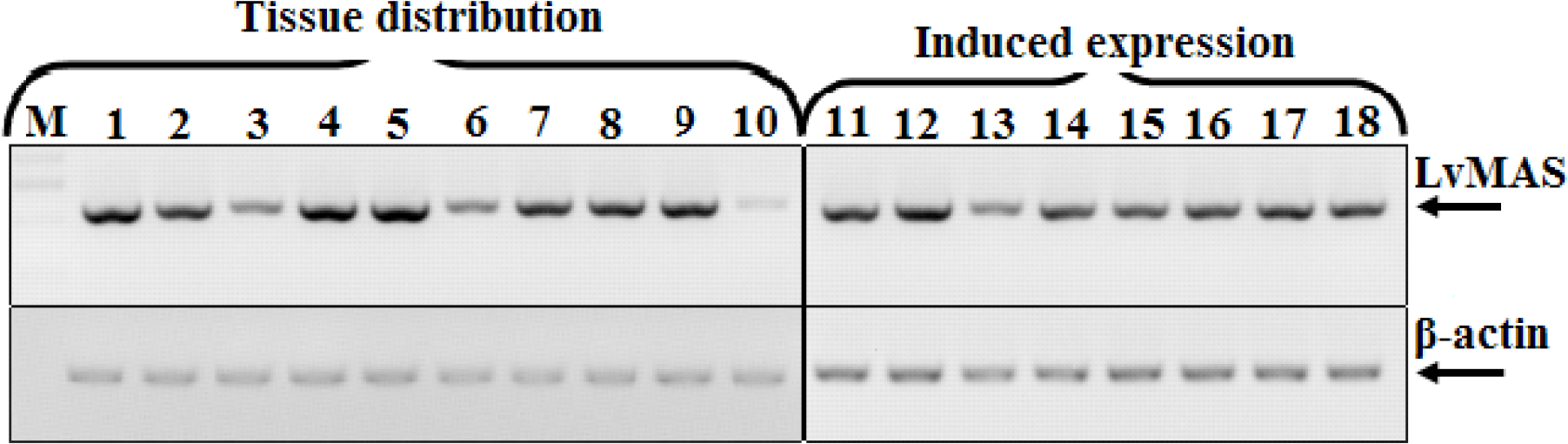
Expression of *LvMAS* mRNA in healthy and immune-challenged shrimps. Tissue distribution of *LvMAS* mRNA in hemocyte (1), epithelium (2), hepatopancreas (3), nerve (4), eyestalk (5), heart (6), pyloric caecum (7), intestine (8), gill (9) and muscle (10) in healthy shrimps by RT-PCR analysis (left). Induced expression of *LvMAS* mRNA in hemocyte by saline (12), LPS (from *E.coli*) (13), Gram-negative *V. alginolyticus* (14), Gram-positive *S. aureus* (15), Yeast *S. cerevisiae* (16), white spot syndrom virus (WSSV) (17) or polycytidylic acid (poly I: C) (18), and untreated hemocyte (12) was used for control (right).

**Fig.3.**
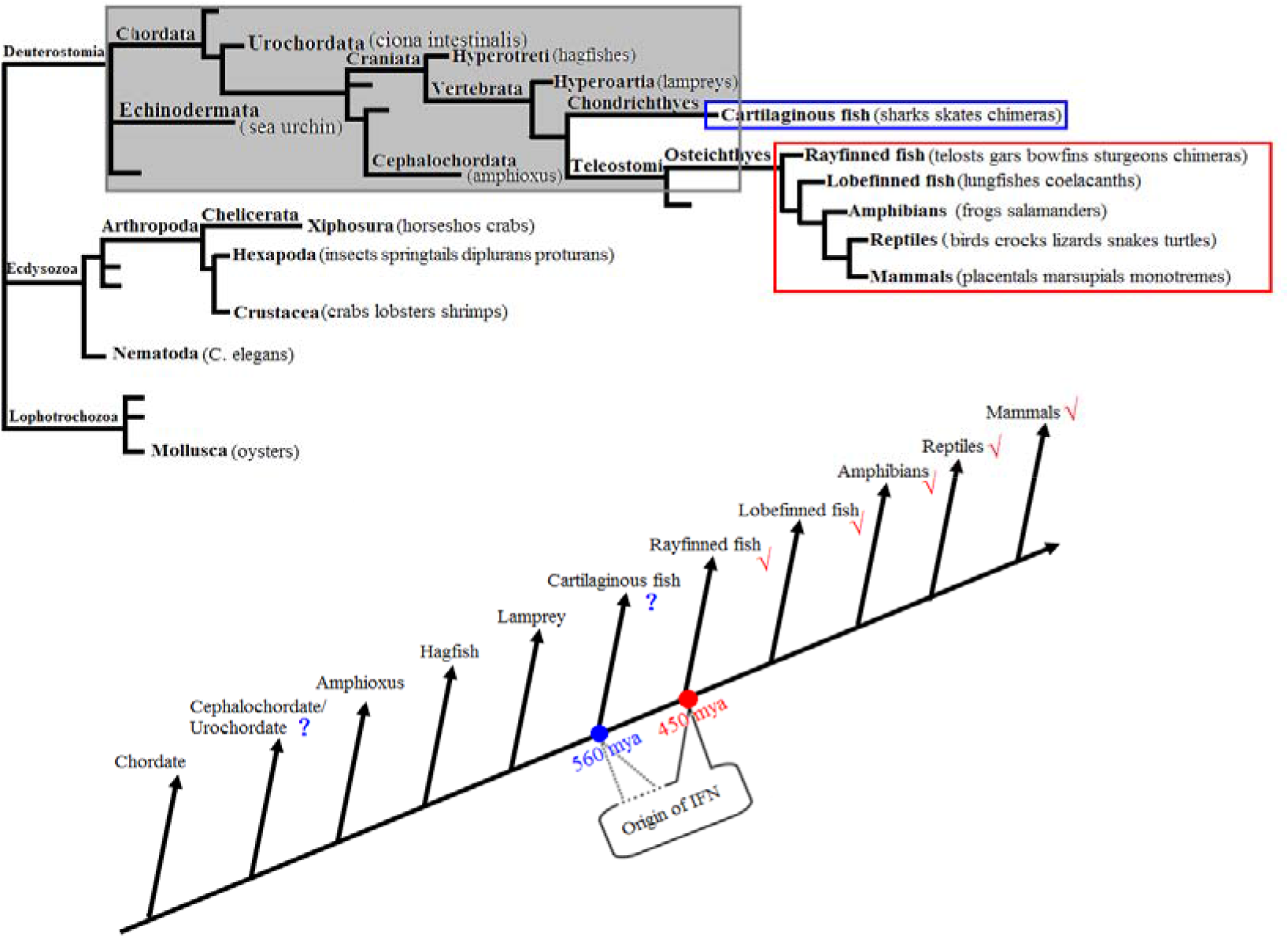
Tree of life showing the emergence and evolution of IFNs.

The encodings of IFNs in animal genomes and EST sequences indicate that IFNs are limited in bony vertebrates (teleost fish, amphibians, reptiles, birds, mammals) and all kinds of interferons are absent in pacific oyster, *C. elegan*, insects, sea urchin *S. purpuratus*, amphioxus, hagfish and lamprey, some of which are consistent with previous studies (Huang et al., 2008; Krause and Pestka, 2005; Savan et al., 2009) (Fig. 3A). Although a human IFN-R1 homology was found in the sea squirt *Ciona intestinalis*, there is no significant IFN-R sequence found in amphioxus, hagfish or lamprey (Krause and Pestka, 2005). Until cartilaginous fish, a shark IFNγ-R is found (Savan et al., 2009). In evolution before cartilaginous fish, most of the vertebrate cytokines except for the tumor necrosis factor like gene are absent, including most interleukins, all interferons, chemokines, colony-stimulating factors, and their cognate receptors (Huang et al., 2008; Krause and Pestka, 2005). The same situation is also observed in other invertebrates including pacific oyster, nematode caenorhabditis elegans, crustacea and genome available insects (Rosa and Barracco, 2008). But from cartilaginous fish some cytokines begin to emerge (Krause and Pestka, 2005; Savan et al., 2009). We propose that vertebrate cytokines especial IFNs origin from cartilaginous fish (Fig. 3B). So we would not agree with the existence of an unknown cytokines to perform the antiviral protection in shrimps (Rosa and Barracco, 2008). In invertebrates such as nematode and insects, RNAi is critical for protecting from viral infections (Saleh et al., 2009; Schott et al., 2005). But RNAi is replaced by the IFNs system in high vertebrates as the primary antiviral responses (Cullen, 2006; Myles et al., 2008). RNAi now exists in vertebrates only as a mechanism of post-transcriptional regulation ‘programmed’ by endogenously encoded miRNA. But RNAi is a sequence-specific gene-silencing mechanism, which are different from the dsRNA-sequence independent unspecific antiviral responses of shrimps (Rosa and Barracco, 2008). We believe that some virus-induced immune proteins such as C-type lectins, hemocyanins and AMPs would play a very important role in this unspecific antiviral responses rather than unknown interferon-like proteins or cytokines similar to vertebrates proposed by Rosa et al (Lei et al., 2008; Zhao et al., 2009). To have a better understanding of shrimp antiviral response, further investigations should focus on these virus-induced immune proteins and a similar RNAi pathway in shrimps. As for the antiviral mechanism of low vertebrates, it is still a gap. Further investigation would be very interesting and contribute to the better understanding of the origin and evolution of animal antiviral system.

In conclusion, this current work demonstrates that IntlP is not a real interferon-like protein, but encoded by a mistaken ORF of MAS. To our knowledge, the reported IFNs are limited in bony vertebrates, and further homology searching and sequence analysis make us believe that most vertebrate cytokines especial IFNs origin from cartilaginous fish.

## Acknowledgements

This research was supported by National Natural Science Foundation of China under grant No. 30325035, National Basic Research Program of China under grand No. 2006CB101802, Chinese National ‘863’ Project under Grant No. 2006AA10A406, 2006AA09Z445, and Foundation from Science and Technology Bureau of Guangdong Province. The nucleotide sequence of LvMAS has been submitted to NCBI database and GenBank with an accession number of DQ923424.

## Abbreviations

IFNs: interferons
MAS: mitochondrial ATP synthase
IntlP: interferon-like protein
ORF: open reading frame
IPTG: isopropyl-β-D-thiogalactopyranoside
LB: Luria broth
LPS: Lipopolysaccharide
ORF: open reading frame
WSSV: white spot syndrome virus
SGIV: Singapore grouper iridovirus
RACE: rapid amplification of cDNA end
RT-PCR: Reverse Transcriptase–Polymerase Chain Reaction
S2: *Drosophila* Schneider 2
SDS-PAGE: sodium dodecyl sulfate polyacrylamide gel
ISKNV: infectious spleen and kidney necrosis virus
MFF: mandarin fish fry
poly I: C polyinosinic polycytidylic acid

